# Warming alters cascading effects of a dominant arthropod predator on microbial community composition in the Arctic

**DOI:** 10.1101/2023.08.16.553601

**Authors:** Amanda M. Koltz, Akihiro Koyama, Matthew Wallenstein

## Abstract

Warming is expected to increase abundances of wolf spider, the top predator in soil communities in the Arctic, but we have little understanding on how increased wolf spider density under warmer conditions affects soil microbial structure through trophic cascades. We tested the effects of wolf spider density and warming on bacterial and fungal community structure in litter through a fully factorial mesocosm experiment in Arctic tundra over two summers. Replicated litter bags were deployed at the soil surface and underground in the organic soil profile and collected at 2- and 14-month incubation. The litter samples were analyzed for community structure of bacteria and fungi and mass weight loss. After 2-month incubation, bacterial and fungal community compositions were already structured interactively by the spider density and warming treatments. Such interaction effect was also found in litter microbial community structure as well as litter mass loss rates after 14-month incubation. Our results show that wolf spiders have indirect, cascading effects on microbial community structure but that warming can alter these effects. The non-linear responses of microbial communities and litter decomposition to warming and increased spider density cast uncertainty in predicting structure and function of Arctic terrestrial ecosystem under warmer conditions in the future.

**IMPORTANCE:** This is one of the first studies demonstrating that predator abundances and increased temperature interactively structure litter microbial communities in the Arctic. The Arctic is one of the fastest warming regions due to climate change and contains disproportionately large amounts of soil organic matter, including thick litter which accumulated over the long time because of slow decomposition. The accelerated soil organic matter decomposition due to the rapid warming can cause positive feedback where resulting greenhouse gas emission contribute to further global warming. Since microbial structure can affect decomposition rates of litter, the observed non-linear responses of soil microbial community compositions and litter decomposition rates indicate challenges in predicting Arctic ecosystem responses in the future.

## INTRODUCTION

Biodiversity and ecosystem processes are largely regulated by abiotic factors relative to biotic interactions in the Arctic due to the long, cold winter (1). However, the Arctic is one of the fastest warming regions on the planet, and warming is expected to accelerate even further due to increasing concentrations of atmospheric greenhouse gases from anthropogenic sources (2–4). Under the predicted warmer climate, the relative role of biotic interactions is expected to increase in regulating major ecosystem processes, such as mineralization of soil organic (C).

One biotic group of particular interest in the warming Arctic is soil microbes, which play the primary role in decomposing soil organic matter. In the Arctic, a large quantity of soil organic matter has accumulated due in part to slow decomposition (5) caused by the combination of low temperature (6), poor water drainage (5) and limited nutrient availability for microbial activities (7, 8). As a result, the northern circumpolar permafrost region, including Arctic tundra, contains approximately 50% of the global soil organic C (9). Under warmer Arctic conditions, organic matter decomposition by soil microbes is predicted to accelerate, which could lead to positive feedback for climate warming (10). Developing a better understanding of how soil microbes respond to warming in the Arctic is critical to assess and predict global C dynamics in the future.

The soil microbial community can be structured by warming via three main pathways: 1) direct abiotic effects of temperature, and indirect biotic interactions mediated by 2) plants and 3) higher level consumers. Microbial community structure may not be sensitive to a few degrees of Celsius warming in a short term (∼ a few months) as demonstrated in lab incubations (11, 12) but see (13). On the other hand, microbial communities may be structured by indirect effects mediated by plants which can instantly respond to warming (14). For example, short-term warming stimulates plant growth, leading to increased C input to soils via litter production and root exudates (15, 16). Warming-induced changes in biotic interactions among microbes and soil fauna can result in changes to microbial structure. Soil microbial communities and their ecosystem processes are determined in part through complex biotic interactions with other community members in the habitat (17). Thus, warming has the potential to indirectly influence microbial structure (18) by altering the composition, abundance, or behavior of soil fauna (19, 20) that consume microbes. Likewise, predators that trigger trophic cascades by altering the abundances or behavior of their litter- and soil-dwelling prey could impact microbial communities (21–23). However, the extent to which predators have the potential to influence soil microbial structure – or whether warming could alter indirect predator effects on the microbial community is understudied.

In Arctic ecosystems, wolf spiders are among the most widespread and locally abundant invertebrate predators (24, 25). The activities of these spiders have cascading effects on lower trophic levels in tundra soil and litter communities with consequences for decomposition. For instance, increased wolf spider abundances stimulated litter decomposition (26). However, the effects of wolf spiders on decomposition were found to be reversed under warmer conditions (27). Although microbes could be affected by changing trophic interactions (28), whether warming combined with shifting wolf spider abundances alters soil microbial communities remains an open question. Several lines of evidence suggest wolf spider populations are expected to increase under warmer conditions across the Arctic (29, 30). Given that these population-level changes are likely to alter intraspecific competition (31) and the top-down cascading effects of wolf spiders on their detrital prey and decomposition (27), these predators are an excellent model system to investigate how warming may alter predator-induced trophic cascades on the soil microbial communities.

In this study, we investigate responses by litter-dwelling fungal and bacterial communities to expected variation in wolf spider density and warming in the Arctic tundra. Specifically, we used field mesocosms and open-topped warming chambers to manipulate densities of generalist-feeding wolf spider predators and ambient temperature over two summers in a well-studied area of moist acidic tundra in Northern Alaska. Previous results from this experiment showed that cascading effects of wolf spiders on decomposition were different under ambient temperature vs. experimental warming (27); after 14 months of in situ litter incubation, higher wolf spider densities resulted in more decomposition under ambient temperature but less decomposition under warming (27). The treatment effects on litter decomposition suggest there may have been interactive effects of wolf spiders and warming on the microbial community. Here, we report on the structure of litter-dwelling bacterial and fungal communities after short (2-month) and longer term (14-month) in situ litter incubation. We hypothesize that the spider density and warming treatments interactively structure fungal and bacterial community structure.

## RESULTS

### Litter characteristics

There were no significant effects of the spider density or warming treatments on litter water content at either soil profile in either year (Fig. 1 and Table 1), indicating that any potential indirect effects of the experimental warming on microbial composition or decomposition due to water content was likely small.

**Fig. 1.**
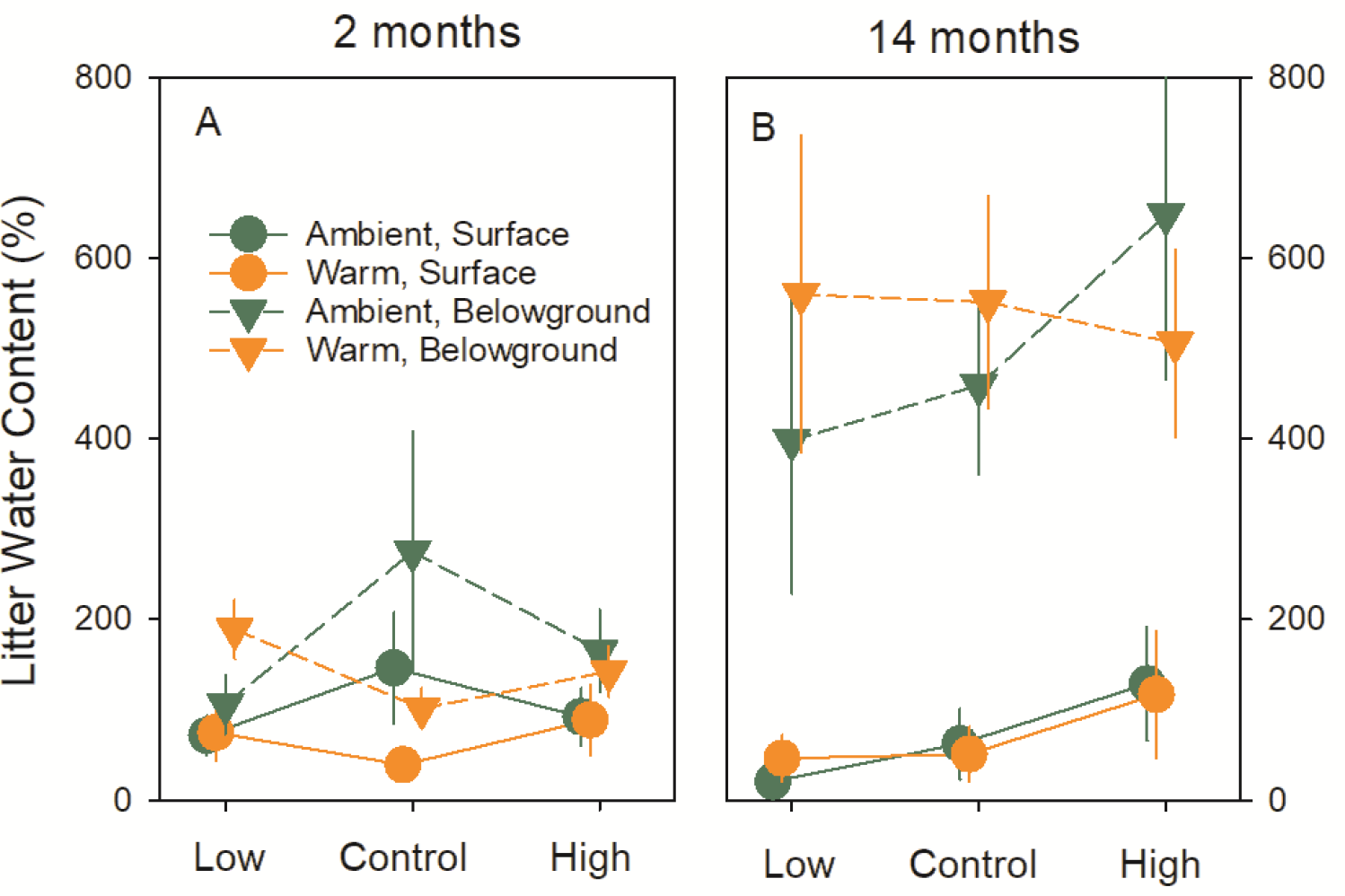
Water content of litter incubated in the field for 2 month (2011) and 14 month (2012). Results of mixed-effect ANOVAs are shown in Table 2. Error bars show standard errors.

**Table 1.** Results of mixed-effect ANOVAs for three litter characteristics (water content, mass remaining and C:N ratio).

| Time | Predictive variables | Litter Water Content |  | Mass Remaining |  | C:N Ratio |  |
| --- | --- | --- | --- | --- | --- | --- | --- |
|  |  | Surface | Belowground | Surface | Belowground | Surface | Belowground |
| 2<br>Months | Spider | $F_{1,20} = 0.43$ | $F_{1,19} = 0.42$ | $F_{1,20} = 1.55$ | $F_{1,19} = 1.52$ | $F_{1,20} = 4.27$ | $F_{1,19} = 0.14$ |
| | | $P = 0.658$ | $P = 0.664$ | $P = 0.236$ | $P = 0.244$ | **$P = 0.029$** | $P = 0.87$ |
| | Temp | $F_{1,20} = 3.28$ | $F_{1,19} = 0.07$ | $F_{1,20} = 1.05$ | $F_{1,19} = 1.09$ | $F_{1,20} = 1.26$ | $F_{1,19} = 0.93$ |
| | | $P = 0.085$ | $P = 0.792$ | $P = 0.317$ | $P = 0.310$ | $P = 0.276$ | $P = 0.347$ |
| | Spider×Temp | $F_{1,20} = 1.3$ | $F_{1,19} = 1.97$ | $F_{1,20} = 2.93$ | $F_{1,19} = 0.29$ | $F_{1,20} = 1.98$ | $F_{1,19} = 0.54$ |
| | | $P = 0.294$ | $P = 0.168$ | $P = 0.076$ | $P = 0.755$ | $P = 0.165$ | $P = 0.594$ |
| 14<br>Months | Spider | $F_{1,20} = 1.2$ | $F_{1,20} = 0.8$ | $F_{1,20} = 1.42$ | $F_{1,20} = 3.08$ | $F_{1,20} = 1.01$ | $F_{1,20} = 1.2$ |
| | | $P = 0.322$ | $P = 0.464$ | $P = 0.264$ | $P = 0.068$ | $P = 0.381$ | $P = 0.322$ |
| | Temp | $F_{1,20} = 0.01$ | $F_{1,20} = 0.41$ | $F_{1,20} < 0.01$ | $F_{1,20} = 1.93$ | $F_{1,20} = 0.07$ | $F_{1,20} = 0.61$ |
| | | $P = 0.904$ | $P = 0.529$ | $P = 0.987$ | $P = 0.180$ | $P = 0.793$ | $P = 0.442$ |
| | Spider×Temp | $F_{1,20} = 0.25$ | $F_{1,20} = 0.53$ | $F_{1,20} = 0.86$ | $F_{1,20} = 8.66$ | $F_{1,20} = 0.32$ | $F_{1,20} = 4.86$ |
| | | $P = 0.785$ | $P = 0.599$ | $P = 0.437$ | **$P = 0.002$** | $P = 0.730$ | **$P = 0.019$** |

705 Significant effects (i.e., $P < 0.05$ ) are in bold.

**Table 2.** Results of mixed-effect ANOVAs for Shannon Diversity Indices and PERMANOVA for NMDS scores.

| Time | Predictive variables | Shannon Index |  | NMDS |  |
| --- | --- | --- | --- | --- | --- |
|  |  | Bacteria | Fungi | Bacteria | Fungi |
| 2 Months | Spider | $F_{2,49} = 1.00$<br>$P = 0.376$ | $F_{2,49} = 0.19$<br>$P = 0.831$ | $F_{2,53} = 2.6$<br>$P = \mathbf{0.042}$ | $F_{2,53} = 2.6$<br>$P = 0.069$ |
| | Temp | $F_{1,49} = 4.51$<br>$P = \mathbf{0.039}$ | $F_{1,49} = 1.77$<br>$P = 0.189$ | $F_{1,53} = 2.24$<br>$P = 0.103$ | $F_{1,53} = 1.45$<br>$P = 0.209$ |
| | Profile | $F_{1,49} < 0.01$<br>$P = 0.998$ | $F_{1,49} = 0.10$<br>$P = 0.755$ | $F_{1,53} = 27.14$<br>$P < \mathbf{0.001}$ | $F_{1,53} = 1.69$<br>$P = 0.189$ |
| | Spider×Temp | $F_{2,49} = 1.47$<br>$P = 0.239$ | $F_{2,49} = 0.67$<br>$P = 0.519$ | $F_{2,53} = 3.14$<br>$P = \mathbf{0.032}$ | $F_{2,53} = 3.82$<br>$P = \mathbf{0.020}$ |
| 14 Months | Spider | $F_{2,38} = 0.90$<br>$P = 0.415$ | $F_{2,40} = 0.48$<br>$P = 0.620$ | $F_{2,44} = 1.84$<br>$P = 0.117$ | $F_{2,44} = 1.69$<br>$P = 0.161$ |
| | Temp | $F_{1,38} = 0.33$<br>$P = 0.570$ | $F_{1,40} = 0.55$<br>$P = 0.464$ | $F_{1,44} = 1.51$<br>$P = 0.231$ | $F_{1,44} = 1.43$<br>$P = 0.203$ |
| | Profile | $F_{1,38} = 0.21$<br>$P = 0.649$ | $F_{1,40} = 19.20$<br>$P < \mathbf{0.001}$ | $F_{1,44} = 5.83$<br>$P = \mathbf{0.007}$ | $F_{1,44} = 6.20$<br>$P = \mathbf{0.006}$ |
| | Spider×Temp | $F_{2,38} = 4.86$<br>$P = \mathbf{0.013}$ | $F_{2,40} = 3.36$<br>$P = \mathbf{0.045}$ | $F_{2,44} = 2.87$<br>$P = \mathbf{0.042}$ | $F_{2,44} = 2.87$<br>$P = \mathbf{0.047}$ |

Reduced models were employed by dropping non-significant higher interaction effects across the analyses. No significant effects were found for Spider×Profile, Temp×Profile or three-way interactions. N.S. indicates no significant effect. Significant effects (i.e., $P < 0.05$ ) are in bold.

The temperature and spider density treatments did not significantly affect remaining litter mass on the surface after 2- or 14-month incubation (Fig. 2A, Table 1). Similarly, there were no significant treatment effects on remaining mass of belowground litter after 2-month incubation (Fig. 2B, Table 1). However, there were significant interactive effects of the temperature and spider density treatments on litter mass after 14-month incubation (*P* = 0.002, Table 1). Specifically, under ambient temperature, the high spider density treatment had less remaining litter mass than the low and control spider density treatments (Fig. 2B). However, under warm temperature, the low spider density treatment had less remaining litter mass than the control and high spider density treatments (Fig. 2B).

**Fig. 2.**
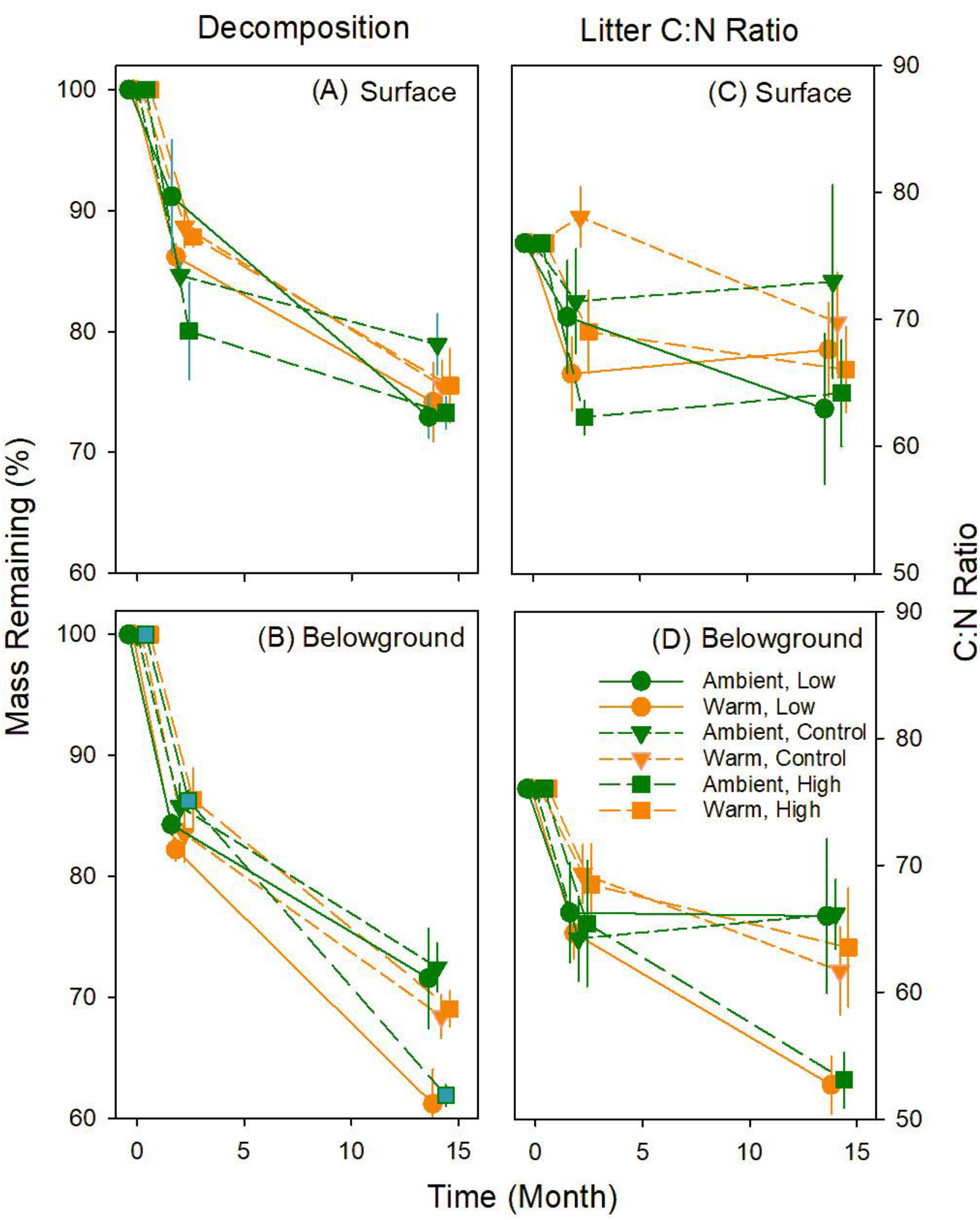
Remaining mass and C:N ratio of *Eriophorum vaginatum* litter recovered after two and 14 months of incubation in the field. Results of mixed-effect ANOVAs are shown in Table 2. Error bars show standard errors.

In terms of litter nutrient content, the high wolf spider densities tended to reduce the C:N ratio of surface litter compared to the control plots after 2-month incubation within each temperature treatment (*P* = 0.029) but that there were no significant treatment effects on surface litter C:N ratio after 14-month incubation (Fig. 2C, Table 1). For the belowground litter, the treatments did not affect C:N ratio after 2-month incubation (Fig. 2D, Table 1) but there were significant interactive effects of the spider density and warming treatments on litter C:N ratio after 14-month incubation (*P* = 0.019, Table 2, Fig. 2D) that reflected the patterns in decomposition (Fig. 2B). Specifically, under ambient temperature, belowground litter C:N ratio was lower in the high spider density than in the low and control spider densities (Fig. 2D).

However, under warm temperature, litter C:N ratio was lower in the low spider density than control and high spider densities (Fig. 2D).

### Alpha diversity of bacteria and fungi

After two months, overall bacterial diversity was lower under the warm than ambient temperature plots (*P* = 0.037, Table 2), a pattern that was most apparent in the higher density wolf spider treatments (Fig. 3A). After 14 months, bacterial diversity was driven by interactive effects of the wolf spider density and warming treatments (*P* = 0.013, Table 2, Fig. 3B). Specifically, in ambient temperature plots, bacterial diversity tended to be higher with increasing spider density, whereas under warmer conditions, bacterial diversity was reduced under higher spider densities at the surface and belowground (Fig. 3B).

**Fig. 3.**
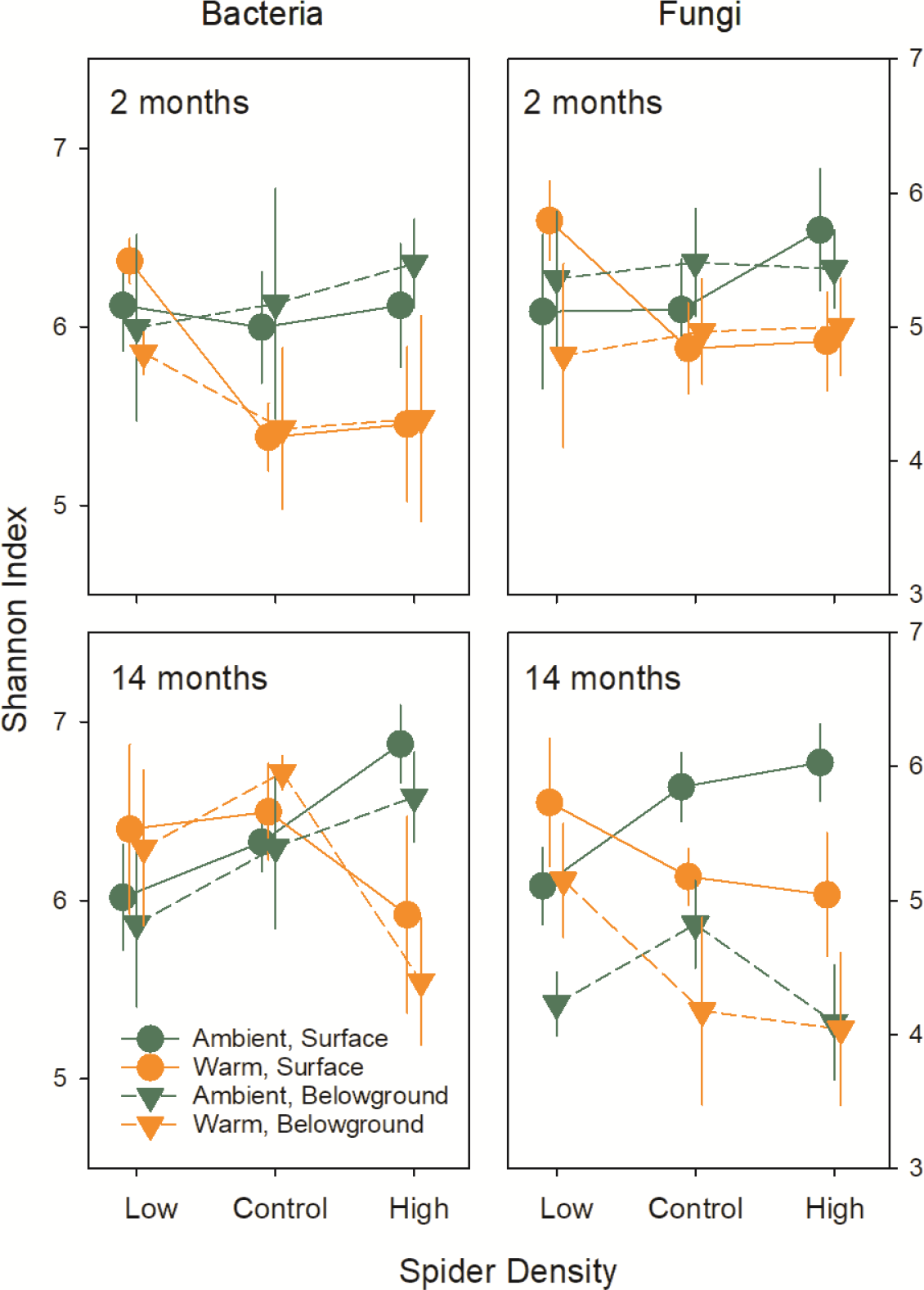
Shannon diversity indices of bacterial and fungal communities in surface and belowground litter collected in 2011 and 2012. Results of mixed-effect ANOVAs are shown in Table 2. Error bars show standard errors.

After the 2-month incubation, fungal diversity was not affected by the experimental wolf spider or warming treatments at either soil profile location (Table 2, Fig. 3C). However, after 14 months, there were significant interactive treatment effects on fungal diversity (*P* = 0.045, Table 2, Fig. 3D). At the soil surface, fungal diversity increased with higher wolf spider densities under ambient temperature but decreased with higher wolf spider densities under warming (Fig. 3D). Belowground, fungal diversity was reduced in the low and high spider density plots compared to the control plot under ambient temperature, whereas fungal diversity decreased with increasing spider density under warm temperature (Fig. 3D).

### Beta diversity of bacteria and fungi

Bacterial community composition at the OTU level was structured interactively by the wolf spider density and warming treatments after both the two- (*P* = 0.032, Table 2, Fig. 4A) and 14-month (*P* = 0.042, Table 2, Fig. 4B) incubations. Overall bacterial community structure was also different between surface and belowground litter bags during both collection periods (Table 2; Fig. 4AB; two-month: *P* < 0.001, 14-month: *P* = 0.007).

**Fig. 4.**
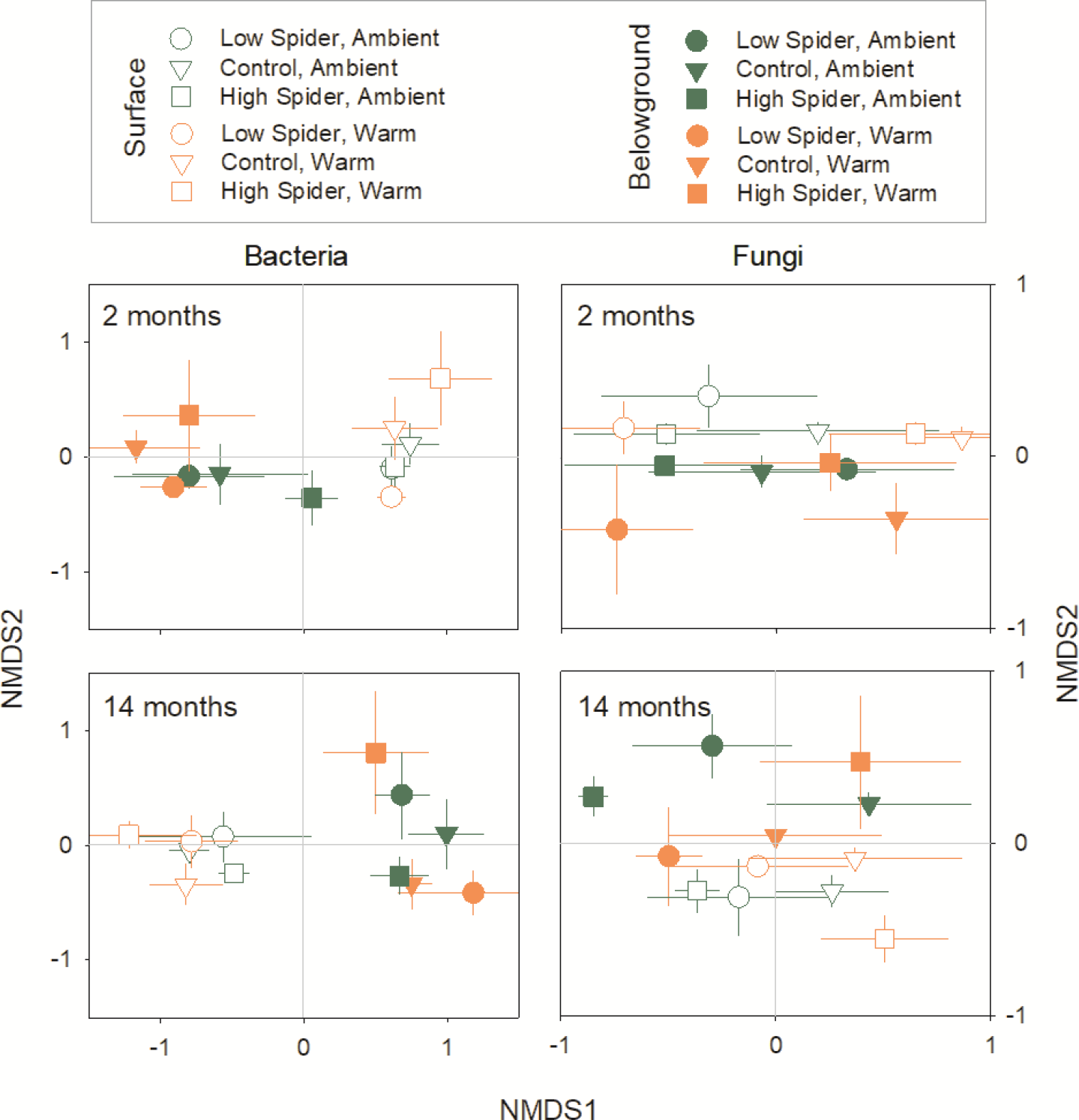
Results of non-metric multi-dimensional scaling (NMDS) for bacterial and fungal communities in surface and belowground litter collected after 2- and 14-month incubation. NMDS was conducted for each microbial group and year. Results of PERMANOVA are shown in Table 2. Error bars show standard errors.

Similar to the bacterial communities, fungal community composition at the OTU level was structured interactively by wolf spider density and the warming treatment after both the shorter two-month (*P* = 0.031, Table 2, Fig. 4C) and longer 14-month incubation periods (*P* = 0.047, Table 2, Fig. 4D). Fungal community structure did not vary between the surface and belowground litter after two months (*P* = 0.189, Table 2, Fig. 4C), but after 14 months, communities were significantly different depending on soil profile (*P* = 0.006, Table 2, Fig. 4D).

### Bacterial and fungal community structure

Overall, Proteobacteria was the most abundant bacterial phylum across the treatments and profiles of litter samples collected from the experimental mesocosms, followed by Acidobacteria, Actinobacteria and Bacteroidetes (Fig. 5). After 2-month incubation, within Proteobacteria, α-Proteobacteria was the dominant class in the surface litter (56.5%), followed by β-Proteobacteria (18.5%), γ-Proteobacteria (3.2%) and δ-Proteobacteria (1.0%) (Fig. 5). However, in the belowground litter, after 2 months, α-and β-Proteobacteria were similarly dominant (33.2% and 35.9%, respectively), and overall γ-Proteobacteria relative abundance was higher compared to the surface litter (10.7% vs. 3.2%). Similar differences between the surface and belowground litter communities were found after 14-month incubation: in the surface litter, α-Proteobacteria was dominant (58.1%) followed by β-Proteobacteria (35.6%), γ-Proteobacteria (1.9%) and δ-Proteobacteria (1.6%) within Proteobacteria (Fig. 5); in the belowground litter, α-Proteobacteria were less abundant than β-Proteobacteria in abundances (28.0% and 35.6%, respectively) (Fig. 5). The relative abundance of γ-Proteobacteria in the belowground litter was higher compared to the surface litter (10.4% and 1.9%, respectively) after 14-month incubation (Fig. 5).

**Fig. 5.**
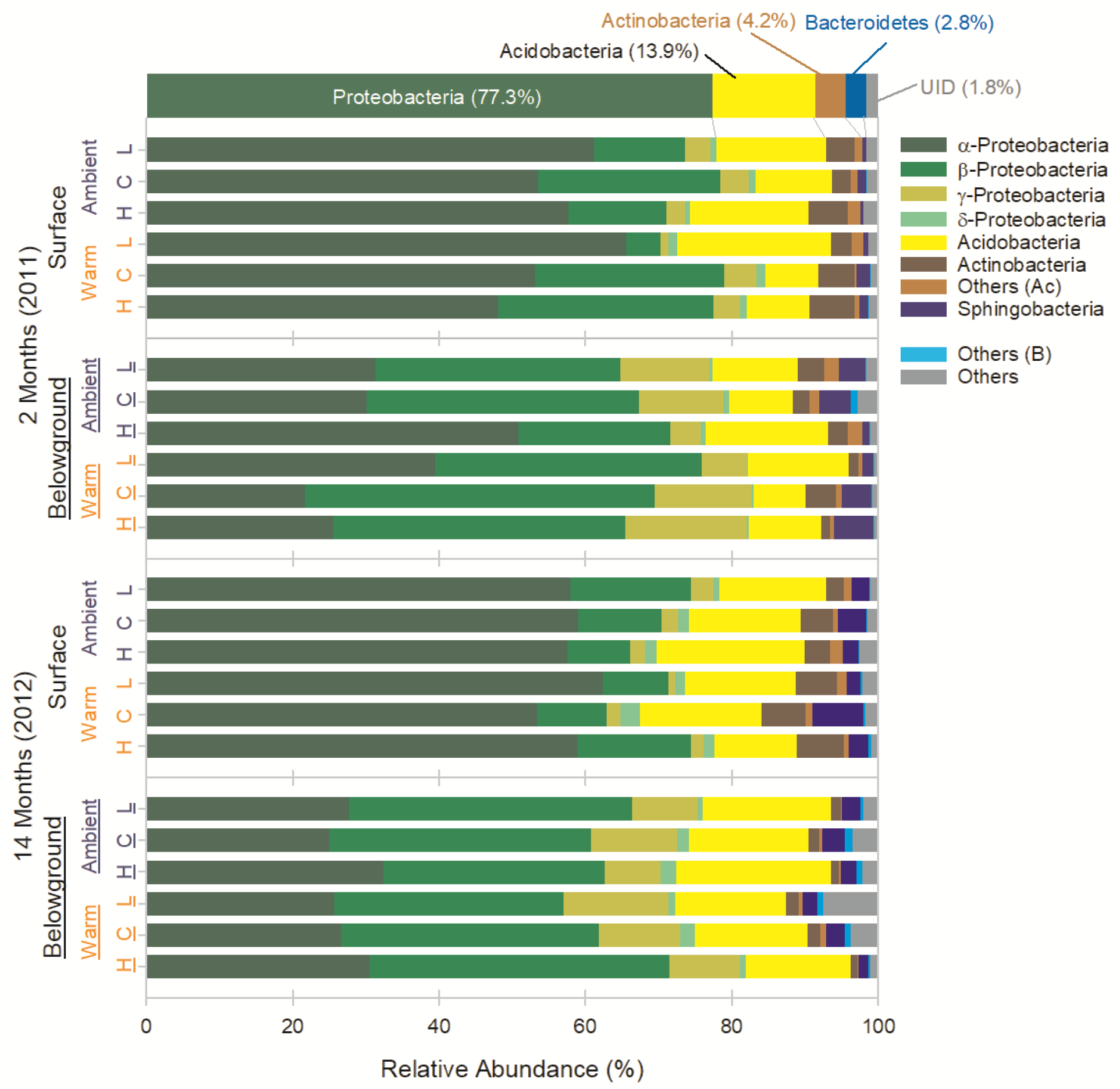
Mean relative abundances of bacteria at the class level in surface and belowground litter collected in 2011 and 2012. Others (Ac) and (B) represent low abundance OTUs belonging to class Actionabacteria and Bacteroidetes, respectively. Others represent OUTs in phyla of low relative abundance other than the four major phyla.

Overall, Ascomycota was the most abundant fungal phylum across the treatments and in both soil profiles, followed by Basidiomycota and Entomophthoromycota (Fig. 6). After 2-month incubation, Leotiomycetes was the dominant class in the surface litter (34.1%), followed by Sordariomycetes (29.7 %), Dothideomycetes (16.8 %) and Saccharomycetes (0.1%) within Ascomycota. In the belowground litter after 2-month incubation, Sordariomycetes were the most abundant (39.9%), followed by Leotimycetes (31.4%), Dothideomycetes (15.0%) and Saccharomycetes (0.9%). Within Basidiomycota, Microbotryomycetes (8.8%) was the dominant class over Agaricomycetes (3.4 %) in the surface litter, whereas, in the belowground litter, Agaricomycetes (7.3%) was more abundant than Microbotryomycetes (0.2%) after 2-month incubation (Fig. 6).

**Fig. 6.**
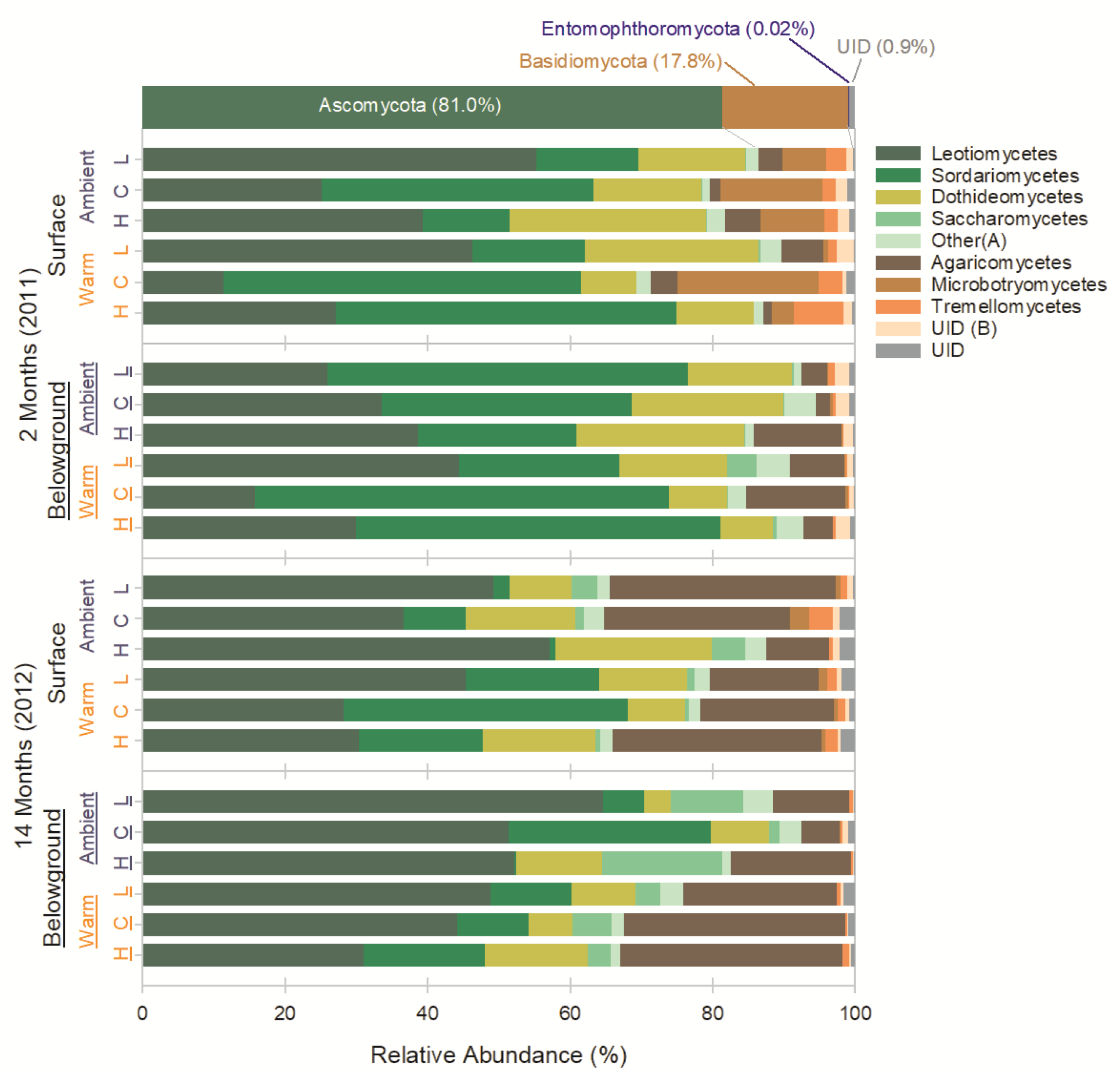
Mean relative abundances of fungi at the class level in surface and belowground litter collected in 2011 and 2012. Others (A) represents low abundance OTUs belonging to class Ascomycota. UID (B) represents OUTs belonging to Basidiomycota and unidentified at the class level. UID represents OTUs unidentified at the phylum level.

After 14-month incubation, Leotiomycetes was the dominant fungal class (40.0%) followed by Sordariomycetes (14.9%) and Dothideomycetes (14.1%), and Saccharomycetes (1.8%) within Ascomycota phylum in the surface litter (Fig. 6). Within phylum Basidiomycota, Agaricomycetes became dominant (22.0%) and relative abundance of Microbotryomycetes (1.0%) and Tremellomycetes (1.6%) in the surface litter (Fig. 6). The belowground litter had similar overall relative abundances to the surface litter after 14-month incubation (Fig. 6): Leotiomycetes was the dominant fungal class (49.2%) followed by Sordariomycetes (12.4%) and Dothideomycetes (8.6%), and Saccharomycetes (6.6%) within Ascomycota phylum, and within Basidiomycota, Agaricomycetes (19.0%) was dominant over Microbotryomycetes (< 0.1%) and Tremellomycetes (0.4%).

## DISCUSSION

### Widespread wolf spider drive changes in litter-dwelling microbial communities and decomposition but these effects change under warming

Previous works have shown that warming (e.g., (32)) and predator abundances (e.g., (18)) can independently structure soil microbial communities. In particular, although studies of indirect effects of predators on microbial communities are rare, the effects of warming on microbial communities have been observed in multiple different terrestrial ecosystems, including Arctic tundra (33, 34), deciduous forests (35, 36), alpine tundra and meadow (37, 38), and grasslands (39–41). However, to our knowledge, the potential for predators and warming to interactively impact microbial structure and function has not been documented before. Notably, we observed effects on microbial communities at soil profile depths below which wolf spiders are active, suggesting that the spatial scale at which spiders influence ecosystem structure and function extends beyond their own microhabitat. Taken together, our findings demonstrate that abiotic conditions and biotic interactions across trophic levels interactively contribute to the structure and function of microbial communities in Arctic tundra. These results provide key insights into the various drivers of non-linear responses of organic matter decomposition in the rapidly warming Arctic, where accumulated litter accounts for a large quantity of mineralizable soil organic C pools (42, 43).

One potential explanation for the interactive effects of spider density and warming on fungal community communities is fungivorous Collembola. Findings from a previously published study from the same experiment demonstrated interactive effects of wolf spiders and warming on lower trophic levels where wolf spiders reduced densities of their fungivorous Collembola prey under ambient temperature, but under warming, effects of higher predatory densities were reversed, causing higher abundances of Collembola (27). There are multiple potential mechanisms behind these interactive effects, including the possibility of intraguild predation among the spider community reducing the strength of top-down predator effects (27). Fungivorous Collembola influence fungal community composition (44, 45) in a variety of ways, including through preferential grazing on particular fungal taxa (46, 47) and by stimulating fungal growth (48). Thus, it is plausible that some of the observed changes in the fungal community were driven by treatment-associated variation in Collembola abundance and grazing pressure.

Shifting species interactions may have also contributed to the observed changes in the litter-dwelling bacterial communities in this experiment. Although the trophic cascade responsible for changes in the bacterial community is unresolved, there are numerous potential interactions that could be affected by wolf spiders. For example, spider density manipulation could affect abundances of bacterivorous nematodes and protozoa through trophic cascades (18). Wolf spiders have been shown to reduce densities of Collembola in a variety of other terrestrial ecosystems (49, 50), suggesting the potential for top-down indirect effects on litter and soil-dwelling microbial communities in those systems as well.

There are several ways through which warming may have influenced microbial community structure in this experiment: 1) direct abiotic effects of warming and/or indirect effects of warming mediated by 2) plants and 3) trophic interactions. Although the relative contributions of these pathways are unknown, our findings and those from other studies suggest the direct effects of warming on microbial composition in the short-term can be small. For example, results from two recent laboratory-based incubation experiments using soils collected from the Arctic tundra showed that microbial community structure was resistant to warming across multiple temperature treatments (11, 12). An Arctic field experiment that increased soil temperature during winter when plants and many arthropods are dormant also showed little changes in soil microbial community compositions for the first 1.5 years (34). Any potential plant-mediated biotic effects on microbial structure were also likely limited by our experimental set-up; surface litter had had little to no contact with plants and the small mesh size on the litter bags may have reduced root-microbial interactions. Therefore, the findings from this experiment indicate that the effects of short-term warming on litter microbial structure were mediated by the indirect effects of wolf spiders and other biotic interactions within the detrital community.

### Interactive effects of wolf spider density and warming were also observed on litter decomposition

However, we do not know how much the observed microbial community compositions uniquely contributed to the decomposition rates in the different treatments. This question is difficult to address due in part to the temporal scale differences; the microbial community compositions were snap shots at given times whereas litter decomposition rates were results of cumulative microbial activities over time. In addition, while fungal traits, such as saprotrophy, can be categorized using fungal taxonomic data bases (e.g., (51, 52)), but it’s challenging to apply it for bacteria which can also play a major role in organic matter decomposition (53). Nevertheless, some studies have demonstrated that microbial structure can influence organic matter decomposition rates: A few controlled lab experiments have shown that manipulated microbial community compositions resulted in different decomposition rates (54–57). For instance, using reciprocal field transplants of inoculated litter across five ecosystems along precipitation and temperature gradients, Glassman et al. (53) demonstrated that abiotic environmental factors were a major predictor for litter decomposition rates, but, to a smaller extent, inoculated microbial community composition and its interaction with the environmental factors accounted for decomposition rates. Thus, it is plausible that the treatment-driven changes in the microbial composition and structure contributed to the observed litter decomposition rates. We note, however, that warming can stimulate soil organic matter decomposition without apparent shifts in soil microbial structure in Arctic ecosystems (58).

### Conclusion

We found that variation in spider density and warming interactively structured litter microbial community composition and modified litter decomposition rates. In polar regions where environmental conditions can be incredibly harsh, abiotic factors are considered the primary drivers of biodiversity and ecosystem processes (59). However, as shown here, widespread arthropod predators can mediate effects of abiotic conditions and therefore alter effects of warming on microbial structure and key ecological processes. Given existing uncertainties around the fate of the soil organic C in the Arctic, the role of predation and other types of biotic interactions in driving ecosystem structure and function warrants further attention as this region continues to warm.

## MATERIALS AND METHODS

### Experimental design

A fully factorial mesocosm field experiment was set up to explore the effects of wolf spider density and warming on microbial community composition and litter decomposition as described in Koltz et al. (27). The experiment was conducted from early June 2011 through late July 2012 near Toolik Field Station (68°38’N and 149°43’W, elevation 760 m) in a well-studied area of moist acidic tundra, which is the dominant tundra type in on the North Slope of Alaska. The average annual temperature is -10°C, with positive temperatures occurring mainly only during the summer months, and the annual precipitation is 200 to 400 mm (60).

A total of thirty plots were randomly assigned to one of six wolf spider density/warming treatments, distributed among five blocks. Half of the plots were warmed using 1.5 meter-diameter ITEX (International Tundra Experiment) open-topped passive warming chambers, which increase mean air temperature by 1 to 2 °C (61). The warming chambers were placed over the plots during June and July of each study year only to avoid affecting snow dynamics. The wolf spider density treatments included: 1) low wolf spider density; 2) control spider density, and 3) enriched wolf spider density. For the low wolf spider density plots, we continuously monitored and removed wolf spiders throughout each summer. Enriched plots received additional spiders in early June of each summer with the aim of bringing wolf spider densities to approximately double the early season average density of the control plots. The efficacy of the wolf spider density treatments (i.e., low, control, and high density) were verified in both years through visual inspection and live pitfall trapping (see (27)).

### Moisture availability

Experimental warming, including through the use of open-topped warming chambers used here, can reduce soil moisture (62) with consequences for microbial community composition (e.g., (41)) and litter decomposition (e.g., 63, 64). To account for this, we measured soil moisture in three locations in each plot at the beginning, middle, and end of the 2012 summer season using a HydroSense portable soil moisture probe (Campbell Scientific, Logan, Utah, USA). Soil moisture data indicated that the warming treatments did not alter average soil moisture content in our experimental plots (*P* = 0.501).

### Litter incubation

Litter bags were used to measure the response of the microbial community to variation in wolf spider densities and to warming. The litter bags were 8 cm by 8 cm with 3 mm mesh size on the top and bottom to allow access by most arthropods (other than wolf spiders and beetles). The bags were filled with 1.5 g of standing dead leaves of the dominant plant, *Eriophorum vaginatum,* which were collected during the previous summer from an area adjacent to our experimental plots, dried at 40 **°**C for 48 hours, mixed, and sub-sampled for litter bag preparation (see (27)). Total C and nitrogen (N) contents were measured for ground subsamples of the initial litter mixture using a CE Elantech Flash EA 1112 Elemental Analyzer (CE Elantech, Inc., Lakewood, New Jersey, USA) at Duke University, Durham, North Carolina, USA.

Two pairs of these litter bags were deployed in each experimental plot during mid-June as described in Koltz et al. (65). From each of these pairs, one litter bag was placed on the soil surface and the other was buried in the litter layer below the moss surface (ca. 5 to 10 cm belowground). One pair of litter bags (i.e., one litter bag from the surface and one from the litter layer) were collected after 2-month incubation and the other pair after 14-month incubation. Upon collection, accumulated soil, ingrown moss and roots, and microarthropods were manually removed from each bag containing decomposed litter and a subsample (0.25 g) of litter was stored at -80 **°**C for DNA extraction at a later date. The remainder of the litter was dried at 40 **°**C for 72 hours to determine litter moisture content and proportional mass loss from the initial litter. Subsamples of dried litter were then ground and analyzed for C and N contents as described above.

### DNA extraction, sequencing of fungal and bacterial communities and sequence data processing

Genomic DNA was extracted from 0.25 g sub-samples of homogenized litter from each collected litter bag using MoBio PowerSoil DNA extraction kit (MO BIO Laboratories, Inc., Carlsbad, California, USA) and eluted genomic DNA samples were stored at -80 °C before downstream processing. The 16S and fungal ITS rRNA genes were amplified for each sample using primer sets of F515F/R806 (66) and ITS1f/ITS2 (67), respectively, which were modified for the Illumina MySeq platform (68).

Polymerase chain reactions (PCR) were performed using triplicate 25-μL assays. Each assay consisted of 12.5 µL of KAPA2G Fast Multiplex Mix (Kapa Biosystems, Woburn, MA, USA), 0.1 µL of BSA (10.0 ng µL^-1^), 1.25 µL of each primer (10.0 µM), and 9.9 µL of a genomic DNA template (1 ng µL^-1^). The PCR thermal cycling steps consisted of an initial denaturation and enzyme activation step of 95 °C for 3 min, followed by 30 cycles of 95 °C for 10 sec, 50 °C for 10 sec and 72 °C for 1 sec. After qualities of PCR products, including amplification and lengths, were assessed by agarose gel electrophoresis, the products were purified using UltraClean® PCR Clean-UP Kit (MO BIO Laboratories, Inc., Carlsbad, California, USA) and quantified using Quant-iT^TM^ PicoGreen® dsDNA Assay Kit (Invitrogen, Molecular Probes, Inc. Eugene, Oregon, USA). Equal quantity of amplicon from each sample was pooled for each of the 16S and fungal ITS PCR products. Each of the pooled amplicons were sequenced with a single run of 2250 bp V2 500-cycle kit on an Illumina MiSeq instrument with at Research Technology Support Facility, Michigan State University, East Lansing, Michigan, USA. All the sequences were deposited at GenBank of the National Center for Biotechnology Information (BioProject ID: PRJNA565353).

Bacterial 16S and fungal ITS Illumina amplicon sequences were processed via the QIIME 1.9.1 toolkit (69). For the fungal ITS sequences, only reverse reads were used for subsequent analyses because some forward reads had poor sequence quality, which would result in substantial reduction in the sequence number per sample in rarefaction. Chimeric sequences in the sequences were identified using USEARCH (70); for the 16S and ITS sequences, the reference-based method with the Greengenes database (version 13.8) (71) and the abundance-based method were used, respectively. The chimeric sequences were removed for the downstream analyses. For the 16S and IST sequences, operational taxonomic units (OTUs) were determined at the 97% similarity level (72) via USEARCH (70) using the Greengenes (13_8 version) (71) and UNITE data base (Version 7) (73), respectively. All the non-bacterial sequences and singletons were removed and rarefied at 22,200 sequences per sample. The remaining sequences were aligned via PyNAST (74) and a bacterial phylogenetic tree was built using FastTree (75).

### Statistical analyses

Linear mixed effects models were used to test the potential interactive effects of our treatments (wolf spider density × warming) and soil profiles (i.e., soil surface or belowground) on the alpha diversity (Shannon Index) of the fungal and bacterial communities. The interaction between spider density and warming and the soil profile of the litter bag were included as fixed effects in the models; experimental block was included as a random effect. Treatment and soil profile effects were estimated separately for fungi and bacteria for each incubation period (2 or 14 months in 2011 and 2012, respectively). Using the same model structure, the effects of treatment and soil profile on the water and nutrient contents (C and N) of the litter contained within the litter bags were also considered. All analyses were conducted using the lmer function of the nlme package (75, 76) in R 3.5.2 (77).

In addition, variation in bacterial and fungal community composition (i.e., beta diversity) from each year were assessed using non-metric multi-dimensional scaling (NMDS, Kruskal 1964) using the *metaMDS* function in the *vegan* package (78) in R. Each model employed two dimensions (*k* = 2) and had an acceptable stress value of < 0.2 (79) (stress = 0.16 and 0.13 for bacteria in 2011 and 2012, respectively, and 0.09 and 0.13 for fungi in 2011 and 2012, respectively). To assess effects of the spider density, warming treatments and soil profiles on microbial composition, permutational multivariate ANOVAs (PERMANOVAs) (80) were performed using the NMDS scores for each microbial group and year with the *adonis* function in the *vegan* package (78). For all models, spider density was treated as a categorical variable (low, control, high density) and treatment block was included as a random effect. For each model, the full interactive model was assessed, and then non-significant predictors were eliminated one at a time to reduce the model. All data were archived through the Arctic LTER Data Catalog (http://arc-lter.ecosystems.mbl.edu/data-catalog).

## ACKNOWLEDGEMENTS

Funding for this research was provided to AMK from the US National Science Foundation (grants 1210704 and 1106401), National Geographic Committee on Research and Exploration, Conservation, Research and Education Opportunities International, Alaska Geographic, the Lewis and Clark Fund, and the Arctic Institute of North America. We thank Kiki Contreras, Samantha Walker, Sarah Meierotto, Jennie McLaren, Jason Stuckey, Greg Selby, PolarTREC teachers Nick LaFave and Nell Kemp, and the staff of Toolik Field Station for assistance in the field. We are also grateful to Guy Beresford, Mary Jane Wolff and Rod Simpson for assistance in the lab.

